# Mutational Scanning of α-Synuclein using a Clickable Protein Tag Reveals Determinants of Membrane-Induced Aggregation

**DOI:** 10.64898/2026.04.20.719647

**Authors:** Daeun Noh, Robert W. Newberry

**Affiliations:** Department of Chemistry, The University of Texas at Austin 105 E 24th St. Austin, TX, 78712, USA

## Abstract

The cellular environment plays a critical role in shaping protein conformations, including aggregated states implicated in disease. One challenge to studying this relationship is that most techniques offering high-resolution insight into the nature of these aggregates cannot be deployed in living cells. Systematic mutagenesis presents an opportunity to bridge this gap but requires general and robust methods to detect protein aggregation across large numbers of variants. Here, we use clickable protein tags to generate FRET pairs in situ that can report protein aggregation in high throughput in living cells. We applied this strategy to probe the nature of cellular inclusions of α-synuclein in a popular yeast model. Our results demonstrate that cellular aggregates of α-synuclein in yeast are likely dominated by protein–membrane interactions, making the aggregation pathway in this cellular model very different than in many in vitro experiments. Furthermore, our comprehensive mutational data reveal the molecular determinants of membrane-induced aggregation. For example, residues that control membrane affinity have a profound effect on membrane-induced aggregation both in vitro and in cells. Furthermore, we discovered that glycine residues, particularly in the central region of the protein, act as gatekeepers to reduce membrane-induced aggregation. Mutational scanning with a clickable protein tag therefore provides high-resolution insights into cellular protein aggregates.

## Introduction

Protein aggregation is a ubiquitous feature of folding landscapes and a driver of human disease.^1–3^ For example, aggregation of the abundant neuronal protein α-synuclein is a central event in Parkinson’s disease.^4,5^ Parkinson’s patients aberrantly accumulate aggregated α-synuclein in degenerating neurons,^6^ mutations associated with familial Parkinson’s disease often increase the rate of α-synuclein aggregation,^7–10^ and α-synuclein aggregates are toxic to cell and animal models.^11–13^ Understanding the nature of protein aggregates and the molecular driving forces of aggregation therefore remain central goals in protein science, but these goals are complicated by the remarkable structural plasticity of many aggregation-prone proteins. α-Synuclein, for example, exists physiologically in a dynamic equilibrium between an intrinsically disordered cytosolic state and α-helical conformations associated with lipid membranes.^14^ During aggregation, α-synuclein can adopt a heterogeneous ensemble of conformations that give rise to oligomeric intermediates, which vary in cytotoxicity, before processing to insoluble amyloid fibrils.^13,15–17^ These fibrils come in many distinct polymorphs,^18–20^ and such structural diversity has been linked to differences in disease phenotypes.^21–23^ More recently, α-synuclein has also been shown to undergo liquid–liquid phase separation,^24–27^ which can promote yet another distinct set of aggregates.^24,28,29^

Efforts to dissect this critical process have relied heavily on in vitro approaches, which can offer detailed information on the misfolded state(s) (e.g., assembly size, conformation, materials properties);^2,30,31^ however, these efforts generally sacrifice environmental context (e.g., binding partners, molecular crowding, subcellular compartmentalization), which can have a profound effect on protein misfolding and aggregation. In contrast, experiments in living cells provide that critical biological context, but few techniques are available to probe the nature of protein aggregates in these complex systems with sufficient resolution to understand the nature of the aggregates. Though a few studies have employed in-cell NMR and/or electron tomography to probe protein aggregates in living cells,^32,33^ most reports of cellular protein aggregation are limited to the observation of subcellular puncta,^34^ whose composition is often poorly understood.

One approach to probing cellular protein aggregates is to compare the behavior of different protein variants, which can help test specific molecular hypotheses regarding the nature of the protein aggregates. For example, if a particular residue is hypothesized to mediate a key interface, its mutation should reduce aggregation. However, the complex folding landscape and enigmatic nature of protein aggregates make designing specific mutations challenging. High-throughput mutagenesis has recently emerged as a powerful approach to bridge this experimental gap.^35–43^ By screening comprehensive missense variant libraries for comparative activity, mutational scanning can help identify key regions, structures, and physicochemical properties that drive biological activity, including protein aggregation. For instance, mutational scanning of Aβ42 aggregation showed that the aggregation of this protein is remarkably robust to substitution, with 75% of substitutions maintaining or increasing amyloid propensity; moreover, the pattern of mutational sensitivity was more consistent with the conformations of brain-seeded fibrils than with fibrils formed from spontaneous Aβ42 aggregation in vitro.^35^ In another study, comprehensive mutagenesis of the TDP-43 prion-like domain identified a mutationally sensitive hotspot region, wherein the pattern of sensitizing mutations suggested formation of both a LARKS (low-complexity amyloid-like kinked segment) β-strand and an α-helix that promote aggregation.^36^ More recently, deep mutational scanning of 1,916 IAPP variants in a yeast-based nucleation assay identified structure formation within a core region that might drive nucleation.^37^

Mutational scanning therefore holds great promise for probing cellular aggregates with sufficient resolution to infer molecular detail. Application of mutational scanning requires assessing the effects of thousands of different mutations, which in turn requires measurements of cellular aggregation that are easily parallelized. Though there are many methods to quantify protein aggregation in living cells, including split fluorescent protein-based sensors,^44,45^ FRET/BiFC aggregation reporters,^46–48^ flow cytometric analysis of fluorescent aggregates,^49^ and high-content fluorescence imaging,^50–52^ most of these methods require either low-throughput imaging or fusing the protein of interest to multiple, distinct reporters. For example, in a prototypical FRET experiment, the protein of interest would be separately fused to donor and acceptor fluorescence proteins, which would need to be co-expressed in the same cell. Unfortunately, this experimental design is not easily translated to mutational scanning because the variant library is cloned as a pool. Creating a reporter FRET pair thus requires creating two pools of variants (i.e., one donor and one acceptor) that must be co-transformed. During transformation, however, is it not possible to ensure that a cell takes up DNA that encodes the exact same protein variant fused to both the FRET donor and acceptor. As a result, cells would express multiple different variants whose distinct activities would be difficult to deconvolute.

Though a variety of strategies exist to circumvent this problem, they generally rely on indirectly tracking downstream effects of aggregation, which can introduce confounding variables and/or limit the systems that can be studied. For example, many high-throughput studies of aggregation-prone proteins track changes in cell survival as a proxy for aggregation;^35,36,38,40^ however, cell survival can be affected by diverse factors besides protein aggregation (e.g., perturbation of native protein function, toxicity arising from non-aggregated states).^53–55^ This approach also requires that the protein aggregates are sufficiently toxic to impart a detectable growth defect, which can require additional engineering of the experimental system. For example, studies of Aβ42 aggregation required fusion to DHFR, in which aggregation-prone Aβ42 variants sequester the fused enzyme into insoluble aggregates and impair its catalytic activity, thereby reducing cell viability under methotrexate selection;^35^ however, the intrinsic toxicity of specific variants can confound interpretation of which mutants contribute to DHFR assembly. Transcriptional reporters can circumvent these challenges by coupling protein aggregation to levels of a specific gene product; for example, a transcriptional reporter approach was developed in yeast (yTRAP), in which the solubility and aggregation state of prion-like proteins regulate the activity of a synthetic transcription factor that controls expression of a fluorescent reporter.^56^ However, because these methods operate through changes in gene expression, they are often specific to particular model organisms.

Forster resonance energy transfer (FRET)-based methods offer an opportunity to fill this gap. The short distances ranges that enable FRET (<10 Å) require intimate molecular contact consistent with protein aggregation. Moreover, because FRET is a general photophysical process, it is fundamentally compatible with a variety of biological systems. However, to adapt FRET to mutational scanning, one must generate a FRET pair without introducing multiple distinct tags during molecular cloning. One prominent approach leverages mEos, a fluorescent protein that undergoes irreversible green to red photoconversion upon UV illumination through photochemical backbone cleavage near its chromophore. This approach has been deployed to quantify cellular protein aggregation at scale, including studies of α-synuclein and the Sup35 prion-like domain.^39,57^ We envisioned a complementary approach that capitalizes on clickable protein tags to generate a FRET pair from a single protein variant in situ. Specifically, the protein of interest can be fused to HaloTag, an engineered, self-labeling protein tag derived from a modified haloalkane dehalogenase that forms a covalent bond with synthetic chloroalkane probes.^58,59^ The promiscuity of HaloTag allows for conjugation with multiple different ligands, including small-molecule fluorophores that can create a FRET pair from a single fusion construct. This circumvents the need to deliver different protein reporters to the same cell. Moreover, this approach should be transferable between different cellular contexts. This approach has already been successfully combined with aggregation-sensitive fluorophores for in-cell detection of protein aggregates,^60^ which we seek to generalize to conventional fluorophores and extend to high-throughput screening of protein aggregates in mutational scanning.

In this study, we used this approach to probe cellular aggregation of α-synuclein in a popular yeast model system.^61^ When heterologously expressed in yeast, α-synuclein recapitulates hallmark pathological features observed in human neurons such as membrane binding, vesicle clustering, dose-dependent toxicity and formation of intracellular inclusions.^11,62^ Accordingly, yeast has been widely used to identify genetic and chemical modifiers of α-synuclein toxicity, many of which have subsequently been validated in neuronal and animal models.^63–69^ Despite this wealth of data, the nature of the cellular aggregates themselves remains unclear. Do the aggregates require amyloid formation? Are the aggregates dominated by protein–protein interactions or by protein–membrane interactions? In vitro data suggest that interactions between α-synuclein and lipid membranes can result in both amyloid nucleation and/or vesicle tethering,^70–73^ but it is not clear which might dominate in any given system, including yeast.

## Results and Discussion

### Development and validation of a HaloTag-based cellular aggregation assay

To probe α-synuclein aggregation in living cells, we employed budding yeast (*Saccharomyces cerevisiae*), a genetically tractable system amenable to high-throughput measurement. Although yeast lacks a direct homolog of α-synuclein, it provides a convenient eukaryotic model for investigating cellular consequences of α-synuclein expression, including protein toxicity and aggregation.^61^ In yeast, heterologous expression of α-synuclein results in spontaneous aggregation, which manifests as punctate intracellular inclusions that are associated with endoplasmic reticulum and Golgi-derived vesicles,^74^ consistent with key features of Lewy bodies, which include α-synuclein-enriched clusters of membranous structures.^74–78^ These phenotypic features develop over the course of hours in yeast, and the organism’s rapid growth enables the generation of large cell populations, thereby enabling mutational scanning.

Here, we developed a FRET-based high-throughput assay to quantify α-synuclein aggregation via flow-cytometry. In this system, α-synuclein was fused to HaloTag, and yeast cells expressing the fused proteins were labeled with two HaloTag ligands (Oregon Green [OG; donor]; and tetramethylrhodamine [TMR; acceptor]) which together constitute a robust FRET donor-acceptor pair (Figure 1A). After expressing the α-synuclein–HaloTag fusion in yeast cells and labelling with donor and acceptor fluorophores, most cells exhibit relatively low FRET efficiency (Figure 1B, Figure S1), but a distinct population show increased FRET efficiency, consistent with α-synuclein aggregation (Figure S2). We can therefore exploit the FRET pair generated from HaloTag conjugation in situ to rapidly identify cells with α-synuclein aggregates.

**Figure 1.**
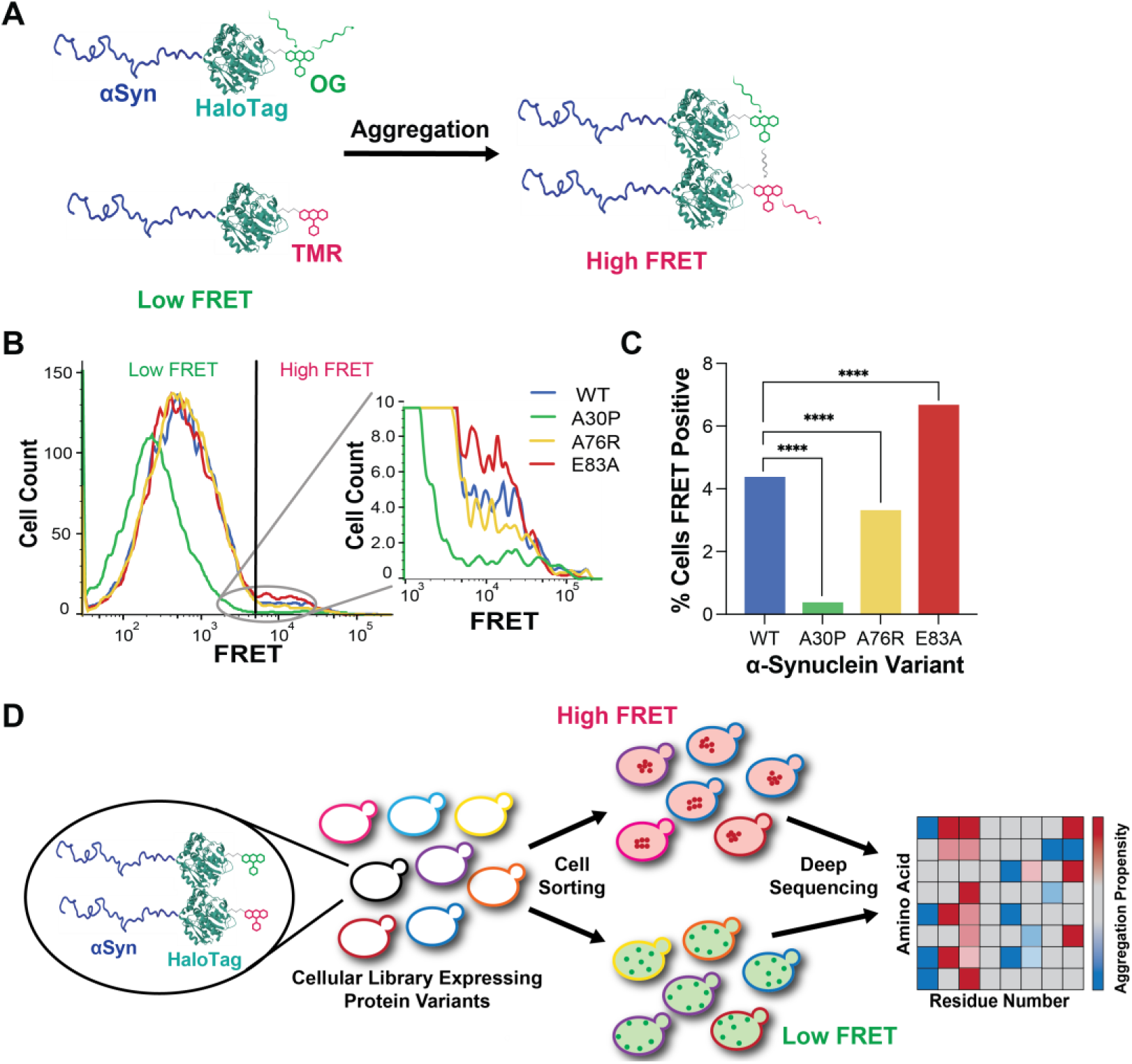
Development of FRET-based assay for mutational scanning of α-synuclein aggregation. (A) Overview of the aggregation assay. α-Synuclein (blue) is genetically fused to HaloTag (teal), which is subsequently labeled with donor and acceptor fluorophores (green and red, respectively) to enable FRET upon α-synuclein aggregation. (B) Distribution of FRET measurements amongst yeast cells expressing α-synuclein variants. (C) Proportion of yeast cell populations exhibiting high-FRET inclusion, as quantified by flow cytometry in B. (n = 10,072 for WT; 10,074 for A30P; 10,072 for A76R; 10,080 for E83A; **** = p<0.0001 by Fisher’s exact test) (D) Overview of the mutational scanning strategy. A comprehensive α-synuclein variant library comprising 2,600 single amino acid substitutions was expressed in yeast. Aggregation is detected through FRET as described in (A). Cells were then sorted based on their FRET signal. The frequencies of each variant within the FRET-negative and FRET-positive populations were quantified by deep sequencing and used to calculate an aggregation score relative to WT.

To validate this assay as a reporter of aggregation, we compared the prevalence of FRET in yeast cells expressing different variants that are known to either reduce (e.g., A30P, A76R) or enhance (e.g., E83A) α-synuclein aggregation in vitro.^79–81^ As expected, both the A30P and A76R mutants dramatically reduced the high-FRET population, whereas the E83A mutation increased the high-FRET population (Figure 1B/C). The HaloTag reporter is therefore sensitive to changes α-synuclein aggregation propensity, allowing us to discriminate variants that both increase and decrease cellular aggregation.

We next extended this approach to deep mutational scanning by constructing a library of α-synuclein missense variants fused to HaloTag (Figure 1D). DNA encoding α-synuclein variants was generated as described previously^82^ and introduced upstream of the HaloTag sequence before transformation into yeast. After labeling with OG and TMR, cells were sorted into low-FRET and high-FRET populations based on their fluorescence profiles (Figure 1D, Figure S3). We sequenced samples collected from two replicate selection experiments. Deep sequencing of the α-synuclein coding region was used to quantify the distribution of α-synuclein variants in low-FRET and high-FRET populations. Aggregation scores were calculated by Enrich2^83^ as the ratio of frequencies of each variant between the two populations normalized to the WT, which were used to derive the sequence–aggregation landscape (Figure 2A).

**Figure 2.**
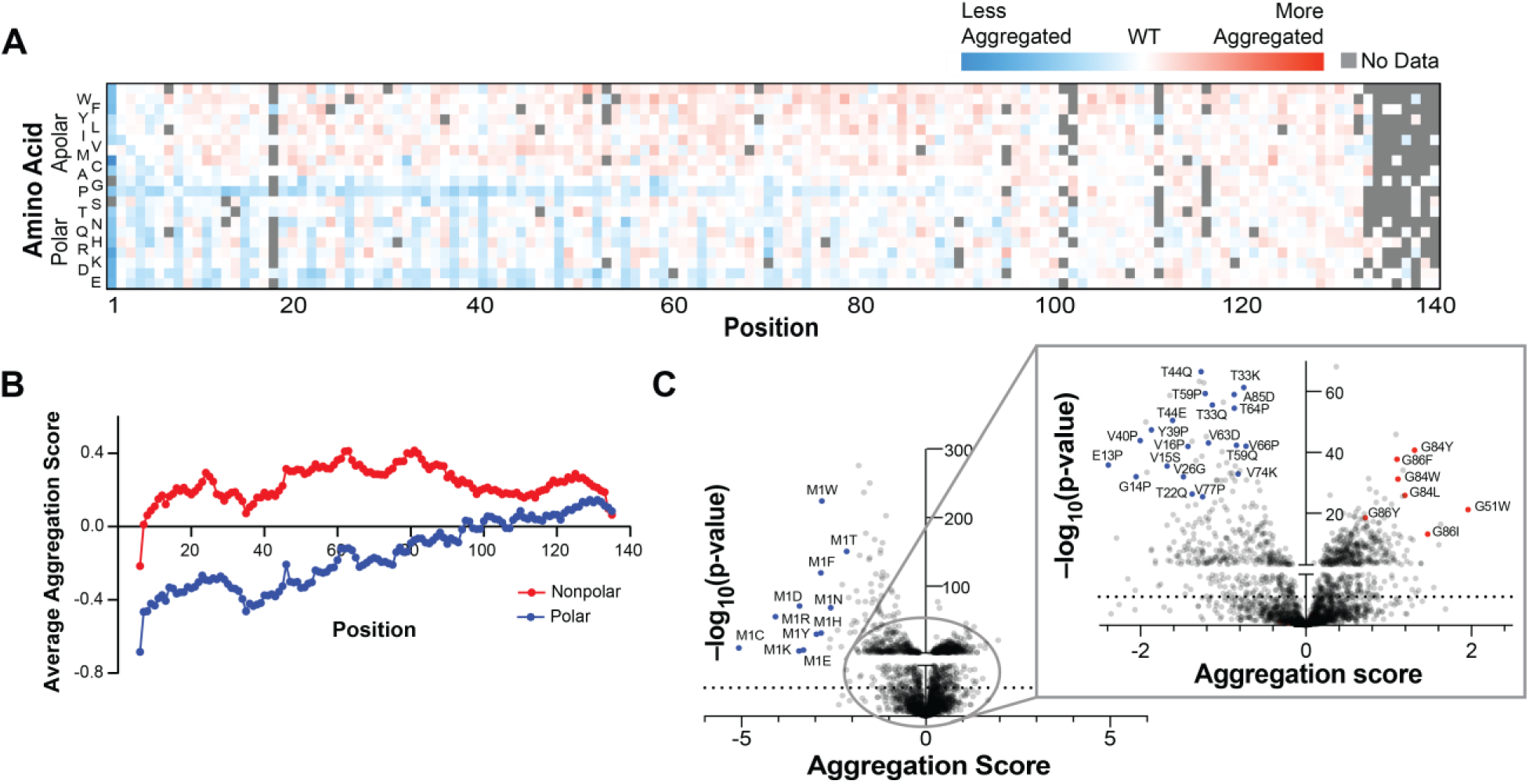
Sequence determinants of α-synuclein aggregation in yeast. (A) Sequence–aggregation landscape of α-synuclein. Aggregation scores were calculated from the ratio of variant frequencies in the FRET-positive versus FRET-negative populations and normalized to WT. (B) Aggregation scores averaged over an 11-residue sliding window for nonpolar (W, F, Y, L, I, V, M, C, A) or polar (S, T, N, Q, H, R, K, D, E) substitutions. (C) Aggregations scores of all single amino acid substitutions in α-synuclein as calculated by Enrich2. The dashed line denotes the 5% false discovery rate (FDR) threshold determined using the Benjamini-Hochberg procedure.

### Sequence–aggregation landscape highlights the central role of membrane binding

As a first insight into the nature of FRET-positive cellular aggregates of α-synuclein in yeast, we calculated aggregation scores for 2,457 single amino-acid substitutions (Figure 2A). Mutations that enhance cellular aggregation were most frequent in the central and C-terminal regions and generally involved incorporating hydrophobic amino acids, which is consistent with the general correlation between protein hydrophobicity and aggregation propensity.^84–86^ When we calculated the average effect of incorporating a hydrophobic residue across the protein (Figure 2B), we observed a maximal effect when hydrophobic residues were incorporated within the non-amyloid component (NAC) region, which is an aggregation-prone segment that commonly forms the core of amyloid fibers. This region might therefore nucleate aberrant interactions, which would likely become more energetically favorable upon increased hydrophobicity. Rather than clustering in any narrow hotspots, incorporation of hydrophobic amino acids across this region increased cellular aggregation, indicating broad participation of these residues in assembly, which is consistent with both the wide variety of known aggregation pathways for α-synuclein,^87–90^ as well as the extensive burial of this region into many different aggregates.^19,91–94^ Though the NAC region is the most hydrophobic region of α-synuclein,^87,95^ its sensitivity to increased hydrophobicity demonstrates that the WT sequence is not fully biased toward aggregation, which could result from selective pressure to avoid aberrant aggregation.

In contrast, mutations that reduce cellular aggregation were more common toward the N terminus and generally involved incorporating additional polar amino acids (Figure 2A). When we calculated the average effect of incorporating a polar residue within different regions of the protein (Figure 2B), we observed that polar amino acids consistently decreased aggregation when incorporated within the first 90-100 residues; however, the strength of aggregation disruption decreased toward the C terminus. Substitutions of the initiator methionine were especially potent in reducing aggregation (Figure 2C), likely due to decreases in protein expression.^82^

We expected that the sequence–aggregation landscape would point to the formation of ordered protein–protein interfaces, such as amyloid; however, several features of the aggregation landscape argue against amyloid formation as the dominant mode of cellular aggregation in this model. For example, amyloid fibrils of α-synuclein adopt cross-β structures in which specific residues form the ordered core (approx. residues 40-90);^21,96^ if amyloid were the dominant FRET-positive species, mutations at these core positions should produce outsized effects on aggregation scores. Instead, sensitive positions were distributed throughout the first ∼100 residues and did not cluster in regions known to form densely packed interfaces. Moreover, the sequence–aggregation landscape did not exhibit the alternating pattern of mutational sensitivity at even and odd positions that would be expected for an in-register parallel β-sheet,^36^ including in the NAC region. We therefore conclude that the dominant FRET-positive species in yeast are not amyloid fibrils, though we cannot exclude minor populations of amyloid or oligomeric species.

Instead, the pattern of mutations that disrupt cellular aggregation demonstrates that FRET-positive α-synuclein aggregates in yeast are dominated by interactions between lipid membranes and α-helical α-synuclein. Proline substitutions were amongst the most disruptive mutations (Figure 2C) and consistently reduced aggregation within the first 90–100 residues (Figure 2A), consistent with proline acting as a helix breaker.^97^ Nonpolar-to-polar substitutions were also amongst the most disruptive (Figure 2C) and recurred every 3-4 residues (Figure 2A), which is consistent with the repetitive structure of the membrane-bound helix. These sensitive positions are predicted to contact the membrane interior when α-synuclein adopts the amphiphilic helix conformation (Figure 3A/B). We therefore expect that the residues that disrupt aggregation reduce the affinity of α-synuclein for lipid bilayers. To test this hypothesis, we compared the effects of mutations on cellular aggregation to the predicted effects of each mutation on membrane-binding affinity. Changes in binding affinity were computed previously by determining the energy associated with placing mutated residues at the appropriate depth within the lipid bilayer, assuming idealized amphiphilic helix structure.^98^ We found strong agreement between computed changes in binding energy and observed aggregation scores (Figure S4; R^2^ =0.56), implicating the membrane-bound helix as a central player in the formation of FRET-positive inclusions in yeast.

**Figure 3.**
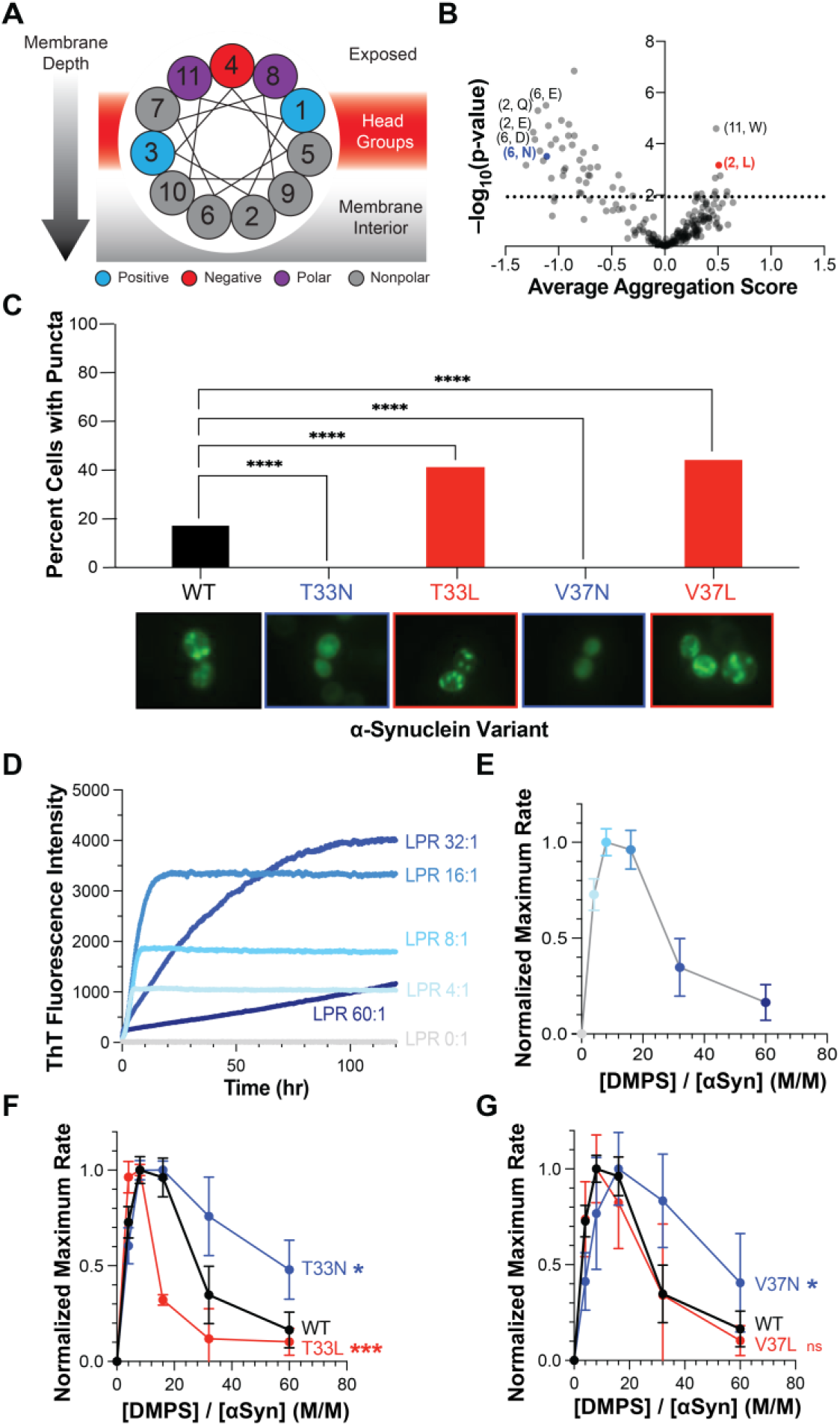
Membrane affinity tunes cellular aggregation and amyloid formation *in vitro*. (A) Helical wheel representation of α-synuclein illustrating the relative orientation of repeated positions in the membrane-binding region. (B) Average aggregation scores of equivalent substitutions at these repeated positions. The dashed line denotes the 5% false discovery rate (FDR) threshold determined using the Benjamini-Hochberg procedure. (C) Aggregation of α-synuclein variants observed by fluorescence microscopy of live yeast (n = 276 for WT; 900 for T33N; 67 for T33L; 136 for V37N; 73 for V37L). (D-E) Rate of α-synuclein amyloid formation in the presence of varying concentration of model membranes (i.e., DMPS SUVs), as observed by thioflavin T fluorescence. (F-G) Normalized maximum lipid-induced aggregation rates of α-synuclein variants with substitutions on the membrane-contacting face of the membrane-bound helix (ns p > 0.05, * p ≤ 0.05, *** p ≤ 0.001, by student’s t-test).

Interactions between α-synuclein and lipid membranes could promote aggregation through multiple mechanisms. First, in vitro experiments show that membrane binding by α-synuclein can nucleate amyloid formation.^70,99^ We found little evidence of this process in the yeast sequence–aggregation landscape, as we did not identify any positions or hotspots whose sensitivity to sterics would suggest a packed interface, nor did we observe the ∼2-residue periodicity expected for β-sheet secondary structure. Another mechanism by which protein–membrane interactions could produce FRET-positive inclusions without requiring amyloid formation is vesicular tethering. α-Synuclein’s amphiphilic helix has been shown to simultaneously engage multiple lipid vesicles,^100–102^ and networks of these interactions could cluster vesicles into higher-order assemblies.^103,104^ This mechanism would be consistent with models developed from solid-state NMR data and electron micrographs of α-synuclein inclusions in yeast,^74^ as well as with α-synuclein’s known physiological function of clustering synaptic vesicles at the presynapse.^105^ Regardless of the precise mechanism by which membrane interactions shape aggregation, these cellular aggregates are distinct from those that are often probed in vitro in the absence of model membranes, underscoring the importance of the cellular environment in shaping the protein conformational landscape. Moreover, distinct cellular environments can promote distinct folding outcomes. For example, when studying seeded α-synuclein aggregation in mammalian cells, Chlebowicz et al. found that mutations in the NAC region consistently reduced aggregation, largely independent of the incorporated residue, pointing to a tightly packed core that is distinct from the cellular aggregates that form in yeast.^39^ The mutational landscape also provided evidence for a locally ordered N-terminal fold that suppresses aggregation by protecting the amyloid-prone core of α-synuclein, which we do not observe in the yeast sequence–aggregation landscape.

### Mutations that affect cellular aggregation also affect membrane-induced amyloid formation

Given that the sequence–aggregation landscape of α-synuclein in yeast appears dominated by the contributions of membrane binding to cellular aggregation, we next asked whether these observations would also shed light on the determinants of other membrane-associated aggregation processes. For example, membrane-binding can nucleate amyloid formation of α-synuclein, in part by concentrating the protein onto a two-dimensional surface, thereby increasing the frequency of aberrant self-associations.^70^ Though we did not observe evidence of amyloid formation in the yeast sequence–aggregation landscape (see above), it is likely operative in other systems.^106–108^ We therefore sought to understand how mutations that affect the cellular aggregation of α-synuclein in yeast contribute to membrane-induced amyloid formation.

Because α-synuclein is a repetitive protein, whose membrane-binding region consists of seven concatenated 11-residue repeats (Figure 3A), we focused on substitutions that caused consistent effects on cellular aggregation across each of the seven repeating segments (Figure 3B). Substitutions that significantly affect cellular aggregation primarily affect positions that are maximally buried into the interior of the lipid bilayer upon binding (i.e., positions 2 and 6 in Figure 3A). For example, incorporating anionic (e.g., aspartate as position 6 or glutamate at positions 2 or 6) or highly polar residues (e.g., asparagine at position 6 or glutamine at position 2) on the hydrophobic face were most effective in reducing cellular aggregation (Figure 3B). In contrast, incorporating a highly hydrophobic residue on the membrane-contacting face (e.g., leucine at position 2) increased cellular aggregation.

To confirm the results of the high-throughput assay, we constructed yeast strains expressing representative α-synuclein variants fused to GFP. Specifically, we converted Thr33 (position 2 in Figure 3A) and Val37 (position 6 in Figure 3A) to either asparagine or leucine, which showed strong effects on cellular aggregation (Figure 3B); asparagine and leucine are also nearly isosteric and neutral in charge, minimizing confounding variables. Consistent with their behavior in the high-throughput assay (Figure 3B), asparagine incorporation at either position 33 or 37 dramatically reduced the frequency of intracellular inclusions, whereas leucine incorporation at those same positions exacerbated cellular aggregation (Figure 3C, Figure S2). Not only do these results support the observations of our high-throughput measurements, but they also highlight the central role of membrane binding in the formation of cellular aggregates of α-synuclein in yeast.

We next hypothesized that these substitutions would similarly affect membrane-induced amyloid formation. To determine if the aggregation determinants observed in yeast apply to membrane-induced amyloid formation, we tracked the fibrillation of purified α-synuclein variants in the presence of model membranes (i.e., small unilamellar vesicles of 1,2-dimyristoyl-sn-*glycero*-3-phospho-L-serine [DMPS]) using the amyloid-binding fluorogenic dye Thioflavin T (ThT).^70,109^ Under quiescent conditions, WT α-synuclein does not form ThT-binding amyloids for at least 100 hours (Figure 3D). Increasing concentrations of DMPS liposomes stimulate increasing rates of amyloid formation due to the presence of surfaces that can promote nucleation (Figure 3D/E). Above a critical [DMPS]:[α-synuclein] ratio (approx. 16:1 for WT), increasing DMPS concentration reduces the rate of amyloid formation, presumably because the increased lipid content dilutes the surface concentration of α-synuclein, reducing the frequency of protein–protein interactions. The relationship between lipid concentration and rate of amyloid formation is therefore inherently biphasic (Figure 3E).^70^

We then compared the membrane-induced amyloid formation of the variants we identified to affect cellular aggregation. Similarly to WT α-synuclein, increasing [DMPS] increased the rate of amyloid formation up to a critical concentration above which increasing [DMPS] slowed amyloid formation (Figure S5), resulting in the expected biphasic relationship (Figure 3F/G). Relative to WT, incorporation of asparagine at either position 33 or 37 increased the concentration of lipid required to maximize the rate of amyloid formation and decreased the sensitivity of the fibrillization rate to further increases in lipid concentration (Figure 3F/G). This behavior is consistent with the polar substitution shifting equilibrium away from the amyloid-inducing, membrane-bound state toward the membrane-unbound state. Indeed, previous titrations of these variants with model membranes demonstrate reduced binding affinity upon asparagine incorporation.^82^ In contrast, substituting Thr33 with leucine decreased the concentration of lipid required to maximize the rate of amyloid formation and increased the sensitivity of the fibrillization rate to further increases in lipid concentration (Figure 3F), consistent with the leucine residue shifting equilibrium toward the membrane-bound state and inducing amyloid formation at lower lipid concentrations. Little effect was seen for the substitution of Val37 with leucine (Figure 3G), consistent with the similar hydrophobicity of these two amino acids. This result highlights the shared role of membrane affinity in both cellular aggregation in yeast, as well as membrane-induced amyloid formation in vitro.

Besides substitutions that affect the polarity of the membrane-contacting face, we also identified a distinct set of substitutions that increase cellular aggregation; specifically, incorporating hydrophobic residues at glycine positions within the NAC domain were some of the strongest enhancers of cellular aggregation (Figure 4A). Intrigued by the prevalence of mutations affecting glycine residues, we plotted the mutational sensitivity of all the glycine residues in the hydrophobic central region of the protein (Figure 4B). Apart from substitutions with proline, a potent helix breaker, or anionic residues that would likely reduce interactions with the phospholipid bilayer, mutations of these glycine residues predominantly increase cellular aggregation. The trend was particularly pronounced at Gly68, Gly84, and Gly86, where even proline and anionic residues caused only weak decreases in cellular aggregation. These observations suggest that glycine residues in the NAC region might act as “gatekeepers” to prevent excessive aggregation, which could contribute to cytotoxicity.^110^

**Figure 4.**
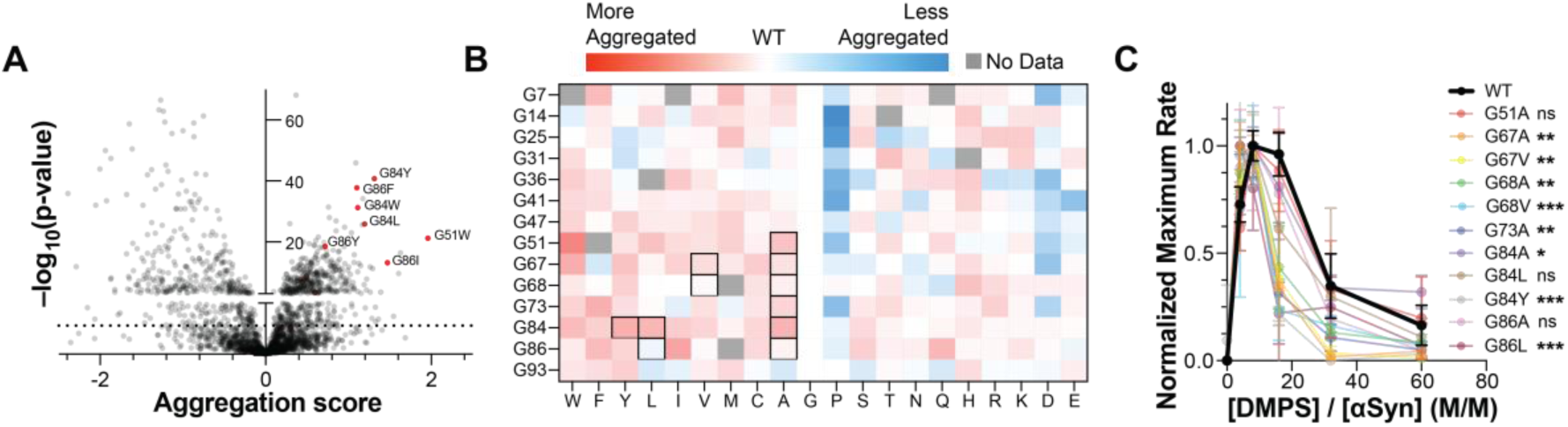
Glycine residues regulate α-synuclein aggregation in cells and *in vitro*. (A) Aggregation scores of single amino-acid substitutions in α-synuclein; the strongest enhancer of aggregation (shown in red) are substitutions of glycine residues with nonpolar residues. (B) Heatmap of aggregation scores for glycine residues in the N-terminal and NAC regions. (C) Normalized maximum lipid-induced aggregation rates of α-synuclein variants with substitutions of glycine residues in the NAC region. ns = p > 0.05, * p ≤ 0.05, ** p ≤ 0.01, *** p ≤ 0.001, by student’s t-test.

To test this hypothesis, we tracked the membrane-induced amyloid formation of eleven different α-synuclein variants in which a glycine within the NAC region had been replaced (Figure 4B/C). We selected a set of variants in which key glycine residues were replaced with either hydrophobic residues, which were some of the most potent aggregation enhancers, or alanine, which also significantly enhanced aggregation (Table S1), despite being the most conservative possible substitution of glycine.^111^ Remarkably, all substitutions tested promoted amyloid formation at lower lipid-to-protein ratios compared to WT, and most did so significantly, including several glycine-to-alanine variants (Figure 4C). Several lines of evidence suggest that these substitutions affect membrane-induced amyloid formation through a mechanism distinct from substitutions that alter the polarity of the membrane-contact face (e.g., Figure 3). First, these residues are positioned at diverse positions around the axis of the amphiphilic helix (Figure S6), and none of them occupy the most buried positions, which are most sensitive to mutation (Figure 2A and 3B). Second, because the membrane-bound helix is more dynamic at its C terminus,^112–114^ residues at the C-terminal end of the helix generally contribute less to binding affinity than residues toward the N terminus.^98^ Moreover, substituting glycine with alanine is predicted to contribute little to binding affinity.^98^ Indeed, when we tested the membrane-binding affinity of representative variants by titrating them with model membranes, we did not observe any significant change in affinity (Figure S6).

We speculate that glycine residues in the NAC region control membrane-induced aggregation by increasing protein dynamics. We initially predicted that replacing glycine residues might reduce aggregation by decreasing dynamics in the membrane-bound state, thereby reducing the probability of engaging in aberrant interactions with either other α-synuclein monomers or nearby lipid bilayers; however, our data are inconsistent with that hypothesis, as replacing glycine residues generally increases aggregation. Instead, we hypothesize that glycine residues contribute to a kinetic barrier aggregation by reducing the probability of adopting an aggregation-competent conformation. Amyloids generally require precise packing, often forming desolvated steric zippers,^115^ so conformational dynamics around glycine residues could reduce the frequency with which the protein samples those conformations. This mechanism could explain why replacing glycine residues in the NAC region cause a decrease in the amount of lipid required to accelerate amyloid formation: as the protein more frequently samples aggregation-competent conformations, each encounter between monomers becomes more likely to lead to amyloid formation, thereby reducing the amount of the membrane-bound nucleus needed to drive fibrillization. For comparison, replacing glycine residues in other functional and pathological amyloid systems, including CsgA and Aβ, similarly increase aggregation propensity.^79,116^ We speculate that a similar mechanism could contribute to the protective effect of glycine residues on cellular aggregation of α-synuclein.

## Conclusions

We find that cellular aggregates of α-synuclein in yeast are fundamentally distinct from those formed in other cell types or in vitro in the absence of model membranes, informing the choice of model system for studying distinct sectors of the conformational landscape. We determined that membrane affinity plays a critical role in the cellular aggregation of α-synuclein in yeast, and we also identify glycine residues as key gatekeepers of membrane-induced aggregation in yeast cells and in vitro. Together, our results highlight the power of mutational scanning for probing the nature and determinants of enigmatic cellular protein aggregates.

## Methods

### α-Synuclein HaloTag Library Construction and Cloning

DNA encoding α-synuclein fused to HaloTag was assembled via Gibson assembly and cloned into the pYES2 expression plasmid (Table S2). DNA encoding α-synuclein variants was obtained from a previously reported plasmid library^82^ and introduced into the pYes2-α-synuclein-HaloTag construct using the EZClone method. Following DpnI digestion, reaction products were column-purified and transformed into *E. coli* TG1 cells by electroporation, yielding approximately 2.65 million transformants. The transformed culture was grown overnight, and plasmids were extracted by miniprep. The plasmid library was bottlenecked to 80,000 members by restrictive transformation into DH5α *E. coli*. The resulting plasmid library was then transformed into *Saccharomyces cerevisiae* strain W303 using the LiAc/PEG/ssDNA method,^117^ generating approximately 0.9 million yeast transformants. Transformed cultures were grown overnight, back-diluted, and returned to log-phase growth. Glycerol stocks were prepared in SCD-URA containing 25% glycerol, with each stock containing ∼10^9^ cells.

### FRET Measurement by Cell Analyzer

α-Synuclein missense variants (A30P, A76R, and E83A) were generated using the QuikChange site-directed mutagenesis methods. All plasmids were confirmed by Sanger sequencing. W303 yeast cells expressing HaloTag-fused α-synuclein WT or point mutants were inoculated from SCD-URA plates into 5 mL SCR-URA medium and grown for 12 h. Cultures were then diluted to at OD 600 of 0.15 in 20 mL SCR-URA and incubated for an additional 1 h. Cells were subsequently diluted again to an OD 600 of 0.15 in 20 mL SCR-URA supplemented with 1% galactose to induce protein expression. After 12 h of induction, 2×10^6^ cells were harvested, washed with PBS, and labeled with HaloTag ligands, Oregon Green (OG) and tetramethylrhodamine (TMR) at 30 °C for 1h with shaking at 225 rpm. The donor dye OG (excitation 496 nm, emission 516 nm) and acceptor dye TMR (excitation 555 nm, emission 585 nm) were used at a 1:5 ratio (2.5 μM OG and 12.5 μM TMR). For flow cytometer compensation, control samples included untreated cells and cells labeled with either donor or acceptor dye alone. To minimize bias from differential cellular uptake of the ligands, OG and TMR were pre-mixed prior to addition to the cells. Following labeling, cells were washed three times with PBS to remove excess dye and resuspended in 1 mL PBS. Flow cytometry analysis was performed using an LSRFortessa flow cytometer (BD biosciences) with the following fluorescence settings: donor channel, 488 nm laser with a 505 nm long-pass and 530/30 band-pass filter; acceptor channel, 561 nm laser with a 582/15 band-pass filter; and FRET channel, 488 nm laser with a 556 nm long-pass and 582/15 band-pass filter.

### Yeast Library Experiments by Cell Sorting

Glycerol stocks of W303 yeast cells carrying the α-synuclein variant library fused to HaloTag were thawed and inoculated into 500 mL SCD-URA. The library cultures were maintained in log phase by dilution into fresh SCD-URA every 12 h during incubation at 30 °C with shaking at 225 rpm. After 12 h, cells were diluted into 0.05 OD in SCR-URA incubated for an additional 12 h at 30 °C. These cells were harvested as uninduced cells and stored at –80 °C. The remaining cells were diluted again to 0.05 OD in SCR-URA supplemented with 1% galactose and incubated for 12 h to induce expression. Cells were collected by centrifugation at 3,000 g for 10 min, washed with PBS, and aliquots corresponding to 20 OD were stored at –80 °C. For labeling and sorting, HaloTag ligands OG and TMR were prepared as 5× concentrated stock solutions. Four labeling conditions were prepared: untreated cells (20 million cells), OG-labeled cells (20 million cells), and TMR-labeled cells (20 million cells), and OG+TMR-labeled cells (100 million cells). Labeling was performed for 1 h at 30 °C, with shaking 250 rpm in the dark. Cells were then washed three times with PBS and resuspended at a concentration of 20 million cells/mL for flow-cytometric sorting. Sorting was performed on a Sony MA 900 cell sorter using a 70 μm nozzle. Compensation was established using untreated cells and single-labeled controls. Fluorescence detection employed the 488 nm laser for OG and the 561 nm laser for TMR, while FRET signals were assessed with the 561 nm laser turned off. Emission was collected using the following filters: OG, 520/50 band-pass; TMR, 585/30 band-pass; and FRET, 582/15 band-pass with 488nm excitation. Target populations were defined as FRET-positive or FRET-negative, and more than 0.6 million cells were collected per group for downstream applications.

### Sample Preparation and Deep Sequencing

Plasmid DNA was isolated from frozen yeast pellets as described previously,^118^ with modifications including the use of the GeneJet plasmid miniprep kit and incubation of pellets in resuspension buffer with the addition of 12.5 μL of 1 M DTT and 10 μL of Zymolyase 20T for 1 h at 37 °C. The α-synuclein coding region was amplified by PCR from plasmid DNA obtained from library samples collected during the sorting experiments. A sequential PCR approach was used to append the Illumina adapters and primer binding sites for sequencing. Amplicons were gel-purified, quantified by Qubit fluorometry, pooled, and sequenced on an Illumina NextSeq 1000 platform in PE 300 mode.

### Calculation of Aggregation Scores

The α-synuclein library was sequenced on an Illumina NextSeq platform using paired-end reads covering the entire coding region. Reads were merged to reconstruct full-length sequences of α-synuclein. Non-overlapping bases were taken directly from reads, and overlapping regions were resolved by consensus. In case of disagreement, the WT base was retained when present in one of the reads. Otherwise, the base with the higher quality score was selected. Merged sequences were analyzed using Enrich2 (v2.0.1), an open-source tool for deep mutational scanning data to quantify the distribution of α-synuclein variants in FRET-positive and FRET-negative populations. Sequencing data from each population (two replicates each) were processed with SeqLib, assigning DNA variants to α-synuclein protein variants. Variants counts were obtained per sample, and enrichment scores were calculated as log ratios of variant frequencies between FRET-positive and FRET-negative populations, normalized to WT. Associated standard errors, p-values, and z-scores for each variant were calculated as implemented in Enrich2. Volcano plots applied a 5% false discovery rate (FDR) threshold by determining the p-value cutoff at which 5% of hits were expected to represent false positives, as determined by the Benjamini-Hochberg procedure.

### Expression and Purification α-synuclein Variants

Plasmids encoding pET28a-α-synuclein (WT and variants) were transformed in *E. coli* BL21 (DE3) cells. Protein expression was induced with 1 mM isopropyl beta-D-1-thiogalactopyranoside (IPTG) at 0.6-0.7 OD. Cell cultures were incubated at 37 °C with shaking at 225 rpm for 3.5 h, and cells were harvested by centrifugation at 4500 rpm for 10 min. Cell pellets were stored at –80 °C until purification of α-synuclein using a previously reported non-chromatographic method.^28,119^ Briefly, the cell pellets were resuspended in lysis buffer (50 mM Tris, 10 mM EDTA and 150 mM NaCl, pH 8) supplemented with 1 mM phenylmethylsulfonyl fluoride (PMSF) and boiled for 10 min. The lysates were centrifuged at 4 °C for 10 min, and the supernatant were collected. Glacial acetic acid (228 μL/mL supernatant) and streptomycin sulphate (136 μL of 10% of solution/ml supernatant) were added, followed by centrifugation at 12,000 g for 10 min at 4 °C. The resulting supernatants were collected, and solid ammonium sulfate was gradually added to 50% saturation over 30-60 min on ice. Precipitated proteins were collected by centrifugation at 12,000 g for 30 min, and the pellets were washed once with 50% ammonium sulfate solution. The washed pellet was resuspended in 700 μL of 100 mM ammonium acetate and precipitated by addition of an equal volume of 100% ethanol. Ethanol precipitation was repeated once, and the proteins were collected by centrifugation. The final pellets were resuspended in 1 mL of Miili-Q water and lyophilized for 1-2 days. Lyophilized proteins were stored at –80 °C until further purification by reverse-phase high-performance liquid chromatograph (RP-HPLC). α-Synuclein variants were loaded onto a reverse-phase C4 column equilibrated with 95% buffer A (0.1% (v/v) trifluoroacetic acid (TFA)) and 5% buffer B (acetonitrile and 0.1% (v/v) TFA) at a flow rate 1 mL/min. Elution was performed using a linear gradient of buffer B from 5 to 90% at a flow rate of 10 mL/min over 30 min. The protein elution was monitored by absorbance at 220 nm and 280 nm. The peaks corresponding to the α-synuclein was collected and analyzed by SDS-PAGE to assess purity.

### Preparation of DMPS Small Unilamellar Vesicles

Dry 1,2-dimyristoyl-sn-*glycero*-3-phospho-L-serine (sodium salt) (DMPS; 840033, Avanti Research) was dissolved in 80% (v/v) chloroform and 20% (v/v) methanol to prepare a 20 mg/mL stock solution. The appropriate volume of DMPS stock solution was aliquoted, and the solvent was removed overnight in a fume hood to form a lipid film. The lipid was then resuspended in 20 mM sodium phosphate (pH 6.5) to a final concentration of 12.5 mM DMPS immediately prior to use. Small unilamellar vesicles (SUVs) of DMPS were prepared by tip sonication (Fisherbrand model 505 sonic dismembrator) at 20% power for 5 min, using a 2s on / 2s off pulse cycle.

### Thioflavin T Aggregation Assay with Lipid

Thioflavin T (ThT) was prepared a 0.5 mM stock solution in 20 mM sodium phosphate (pH 6.5). α-synuclein protein samples were diluted to a final concentration of 50 μM in 20 mM sodium phosphate (pH 6.5), and lipid vesicles were added to achieve the indicated lipid-to-protein molar ratios. ThT was added to a final concentration of 10 μM. Samples were prepared in a half-area 96 well plate (Corning 3881) with a final volume of 100 μL per well. Plates were sealed to prevent evaporation and incubated at 30 °C under quiescent conditions. ThT fluorescence was measured using a BioTek Synergy H1 microplate reader with excitation at 440 nm and emission at 485 nm. Fluorescence measurements were recorded every 10 min over three days or more. All experiments were performed in triplicates. Linear regression was performed on ThT fluorescence data using overlapping sliding 1-h time windows to determine maximum aggregation rates. The maximum aggregate rates determined for each lipid-to-protein molar ratio were normalized relative to the maximum rate observed across conditions.

### Fluorescence Microscopy and Puncta Quantification

W303 yeast strains expressing α-synuclein-EGFP were inoculated into SCR-URA and maintained in log-phase growth for 12 h. Cultures were diluted to 0.2 OD in SCR-URA with 1% galactose. Yeast cells expressing WT α-synuclein and membrane binding affinity variants (T33N/L, V37N/L) were imaged 12 h later using a Nikon Ti-E microscope equipped with a Plan Apo 40x/0.99 Corr air objective. Fluorescence excitation was provided by a Sutter LS-2 xenon arc lamp, and images were acquired using Semrock five-band filter set for FITC filter set (485/20 excitation, 525/30 emission). Images were captured with a DS-Qi2 monochrome camera and processed using NIS-Elements 4.30. Cells displaying α-synuclein puncta and total cell numbers were quantified manually using ImageJ. Statistical analysis was performed using Fisher’s exact test in Graph Pad Prism 9. The puncta of α-synuclein variants at glycine positions were imaged under a Evos FL auto 2 fluorescence microscope using the Invitrogen EVOS Light Cube, GFP 2.0 (482/25 excitation and 524/24 emission) with LWD EVOS 60x objective (fluorite, NA=0.75, WD=2.2 mm) for capturing high magnification image.

## Data availability

Raw sequencing data are available at the NCBI Sequence Read Archive (PRJNA1450116). Counts of missense variants in FRET positive and FRET negative populations for each replicate are available as supporting tables, as are the resulting aggregation scores. The aggregation scores with associated statistics are available in the supporting tables. Aggregation scores are also available at MaveDB (mavedb.org; urn:mavedb:00001272-a). The remaining data underlying this study are available in the article and its supporting information.

## Supporting information

Figure S1. Yeast cells expressing α-synuclein-HaloTag and labeled with donor (OG) and acceptor (TMR) fluorophores exhibit FRET during flow cytometry.

Figure S2. Fluorescence micrographs of W303 cells expressing α-synuclein variants-EGFP. Figure S3. Yeast cells expressing α-synuclein library-HaloTag and labeled with donor (OG) and acceptor (TMR) fluorophores exhibit FRET during sorting experiments.

Figure S4. Correlation between aggregation and predicted change in membrane affinity averaged at each position.

Figure S5. Thioflavin-T aggregation kinetics of α-synuclein variants at different ratios of DMPS lipid-to-protein ratios.

Figure S6. (A) Selected α-synuclein residues with the membrane surface. (B) Circular Dichroism Titration of purified α-synuclein WT and two non-polar variants at glycine positions with DMPS lipid vesicles.

Table S1. Aggregation scores and p-values of selected nonpolar substitutions at glycine residues in the NAC region.

Table S2. List of DNA sequences.

Table S3. Counts of each variant in each FRET-negative and -positive population.

Table S4. Table of aggregation scores.

Table S5. Table of statistical significance by p-values for α-synuclein single missense variants. Table S6. Table of statistical significance by p-values for α-synuclein equivalent variants in the seven repeating 11-residue regions.

## Acknowledgments

This work was supported by funding from NIH (R00-NS116679 and R35-GM160477) to RWN. We gratefully acknowledge the UT Austin Genomic Sequencing and Analysis Facility (GSAF) for next-generation sequencing support and the Center for Biomedical Research Support (CBRS) Microscopy and Flow Cytometry Facility for assistance with flow cytometry and cell sorting.

## For Table of Contents Only

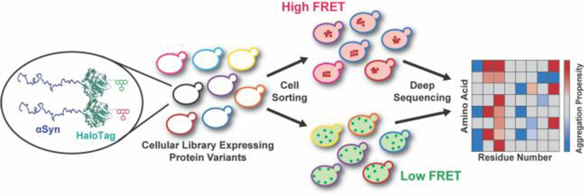

Systematic mutagenesis can probe enigmatic cellular protein aggregates when coupled to an aggregation reporter assay that generates a FRET pair in situ by conjugating fluorophores to a protein tag.

## Notes

### Competing Interest Statement

The authors have declared no competing interest.

