## Supplementary material for "Mutational Scanning of α-Synuclein using a Clickable Protein Tag Reveals Determinants of Membrane-Induced Aggregation": Figure S

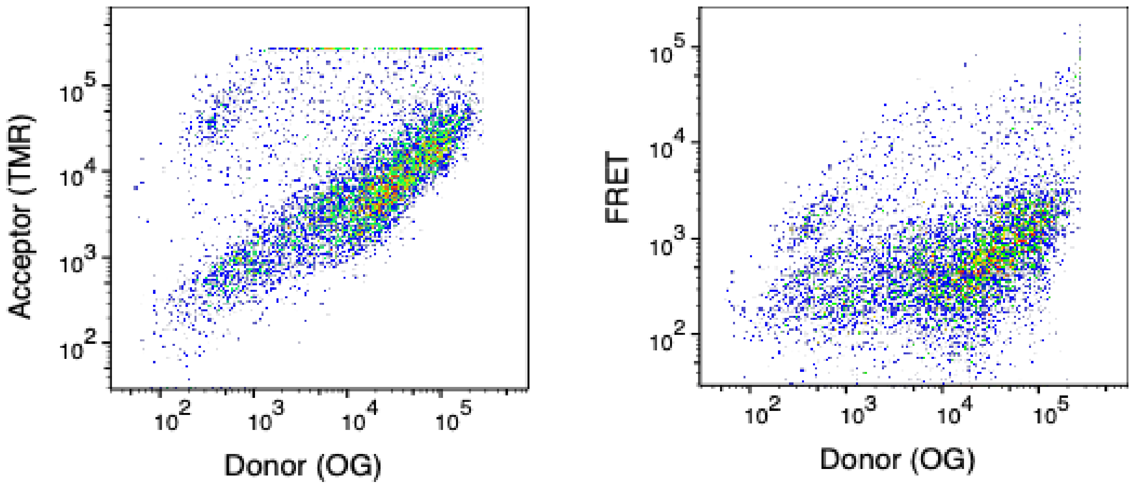


**Figure S1.** Yeast cells expressing α-synuclein-HaloTag and labeled with donor (OG) and acceptor (TMR) fluorophores exhibit FRET during flow cytometry.

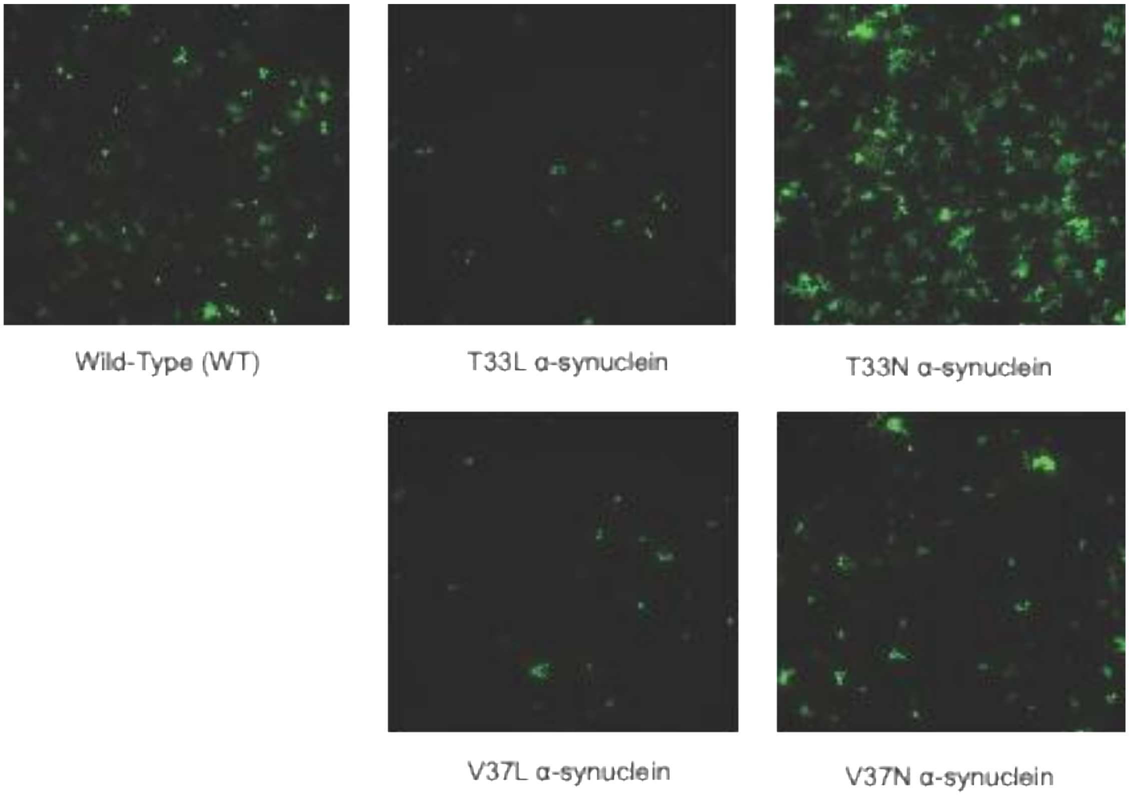


**Figure S2**. Fluorescence micrographs of W303 cells expressing α-synuclein variants-EGFP.

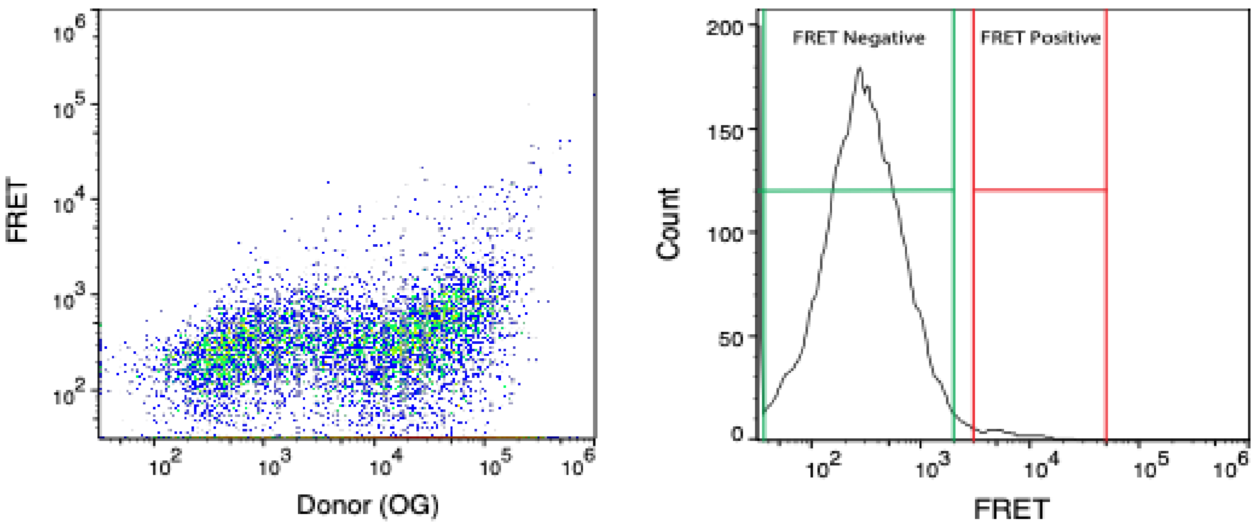


**Figure S3.** Yeast cells expressing α-synuclein library-HaloTag and labeled with donor (OG) and acceptor (TMR) fluorophores exhibit FRET during sorting experiments.

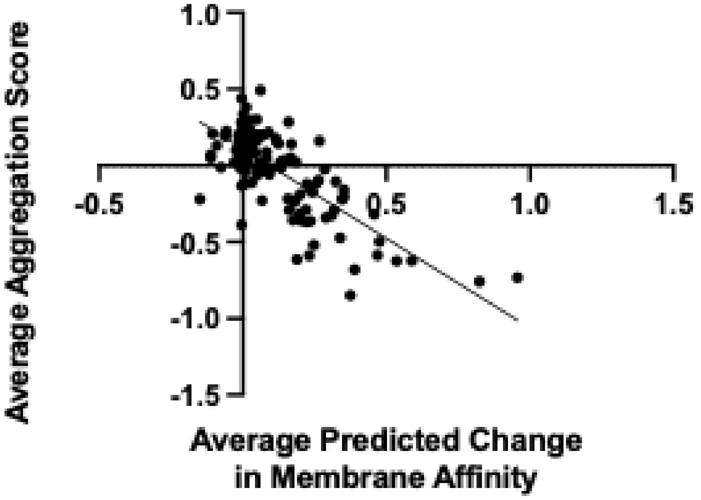


**Figure S4**. Correlation between aggregation and predicted change in membrane affinity averaged at each position. The predicted change data from Newberry et al., (2020), *Nat. Chem. Biol.* were incorporated into the analysis.

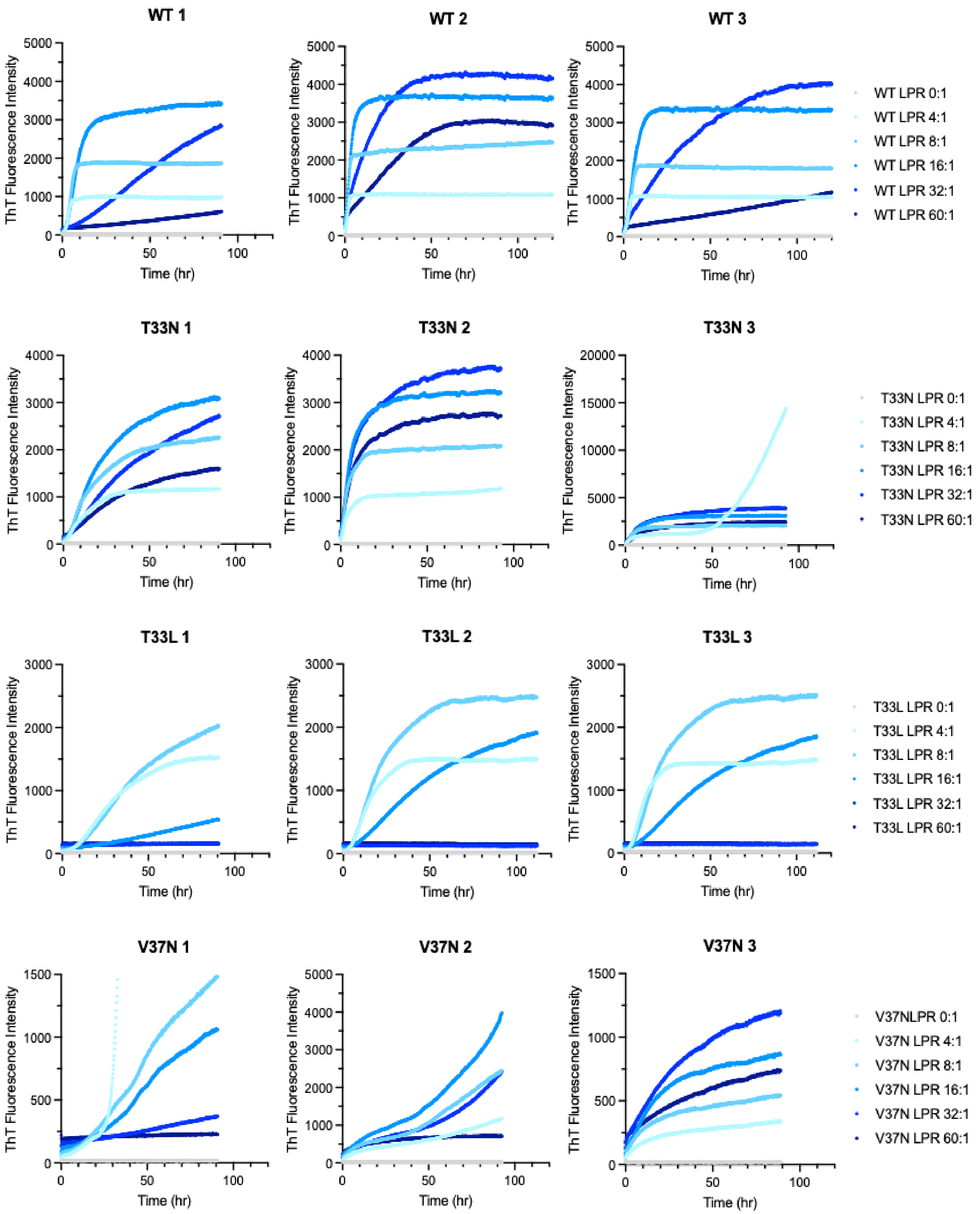
WT 1
WT 2
WT 3
WT LPR 0:1
WT LPR 4:1
WT LPR 8:1
WT LPR 16:1
WT LPR 32:1
WT LPR 60:1
T33N 1
T33N 2
T33N 3
T33N LPR 0:1
T33N LPR 4:1
T33N LPR 8:1
T33N LPR 16:1
T33N LPR 32:1
T33N LPR 60:1
T33L 1
T33L 2
T33L 3
T33L LPR 0:1
T33L LPR 4:1
T33L LPR 8:1
T33L LPR 16:1
T33L LPR 32:1
T33L LPR 60:1
V37N 1
V37N 2
V37N 3
V37NLPR 0:1
V37N LPR 4:1
V37N LPR 8:1
V37N LPR 16:1
V37N LPR 32:1
V37N LPR 60:1
ThT Fluorescence Intensity
Time (hr)

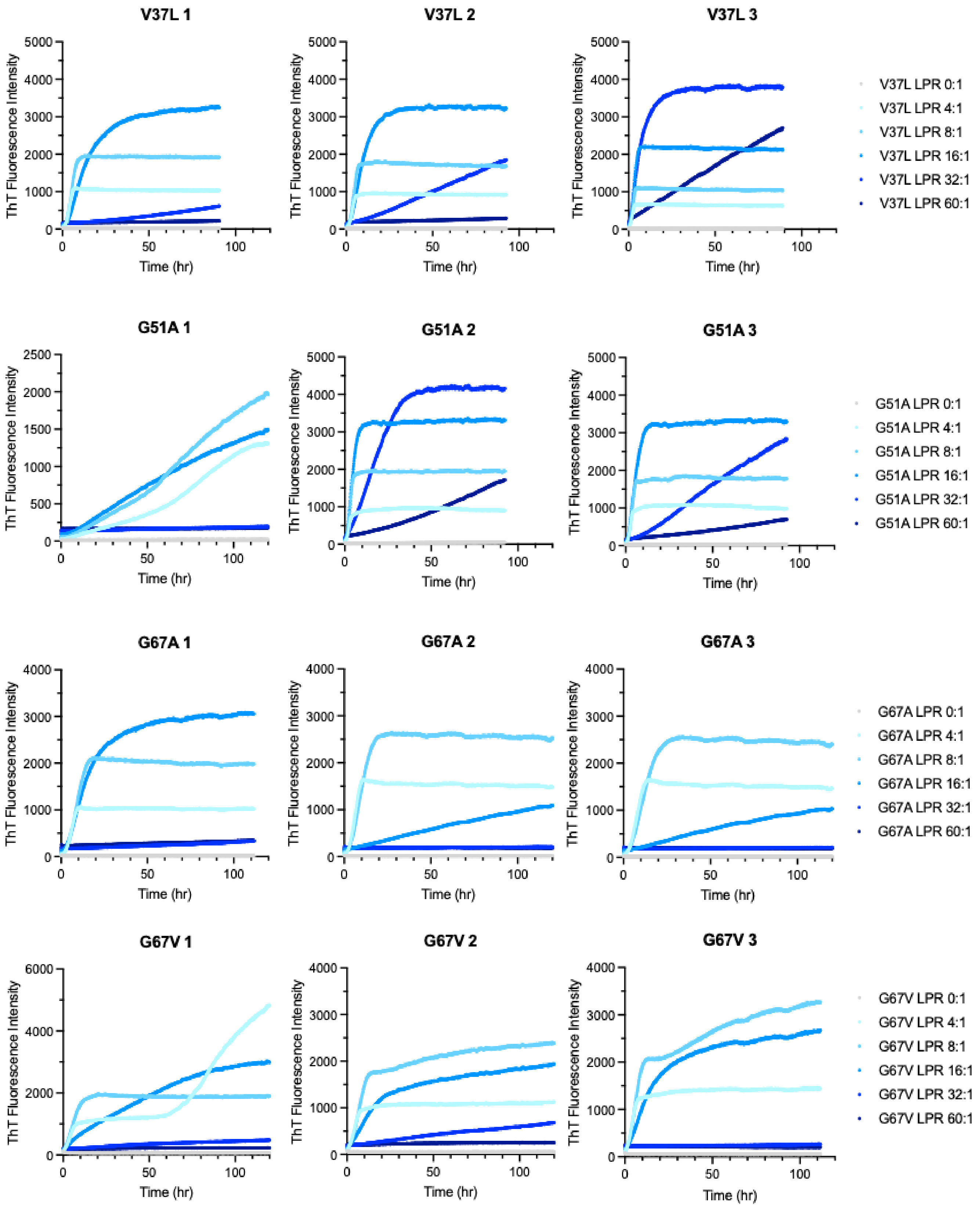
V37L 1
V37L 2
V37L 3
ThT Fluorescence Intensity
Time (hr)
V37L LPR 0:1
V37L LPR 4:1
V37L LPR 8:1
V37L LPR 16:1
V37L LPR 32:1
V37L LPR 60:1
G51A 1
G51A 2
G51A 3
G51A LPR 0:1
G51A LPR 4:1
G51A LPR 8:1
G51A LPR 16:1
G51A LPR 32:1
G51A LPR 60:1
G67A 1
G67A 2
G67A 3
G67A LPR 0:1
G67A LPR 4:1
G67A LPR 8:1
G67A LPR 16:1
G67A LPR 32:1
G67A LPR 60:1
G67V 1
G67V 2
G67V 3
G67V LPR 0:1
G67V LPR 4:1
G67V LPR 8:1
G67V LPR 16:1
G67V LPR 32:1
G67V LPR 60:1

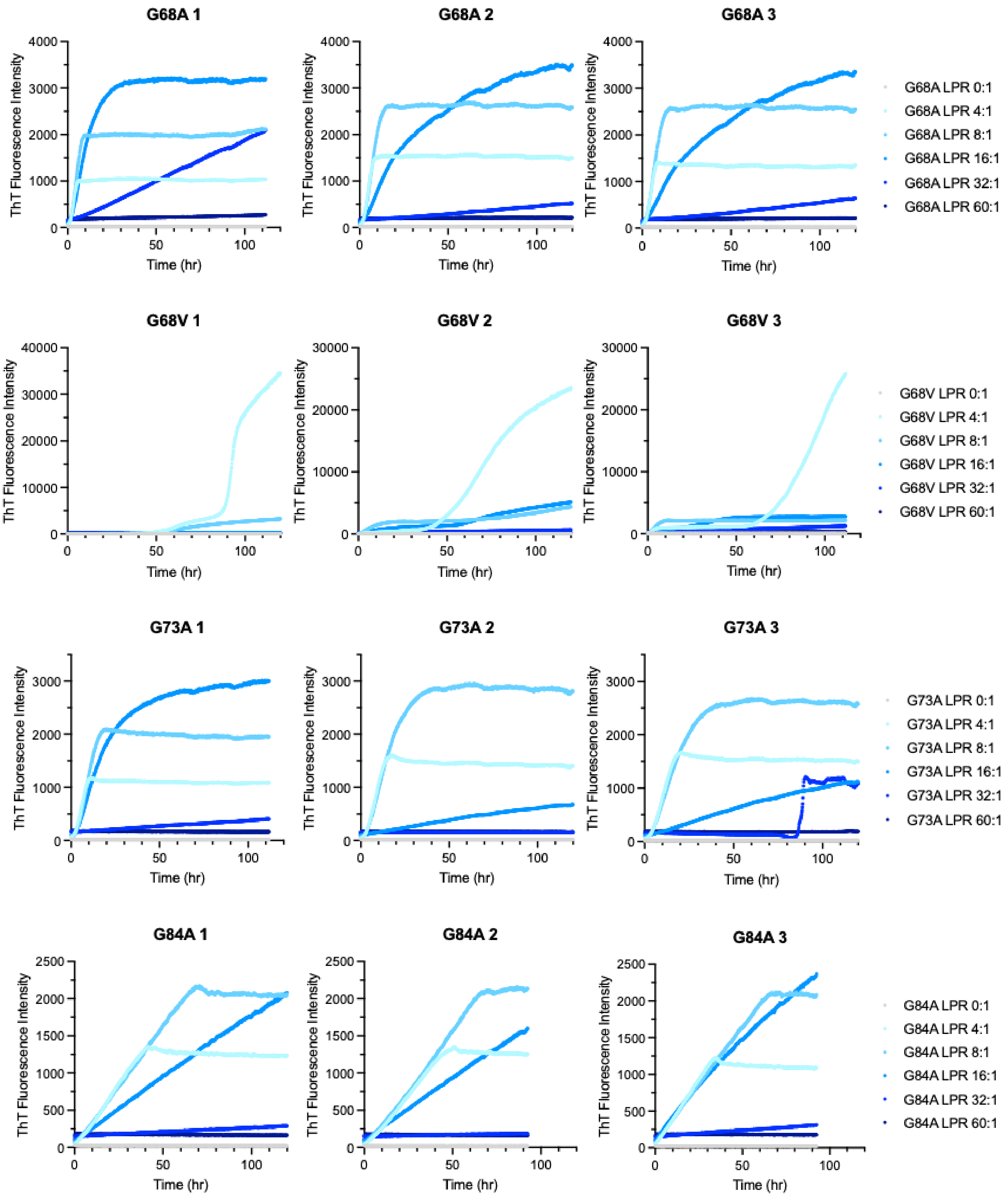
G68A 1
G68A 2
G68A 3
ThT Fluorescence Intensity
Time (hr)
G68A LPR 0:1
G68A LPR 4:1
G68A LPR 8:1
G68A LPR 16:1
G68A LPR 32:1
G68A LPR 60:1
G68V 1
G68V 2
G68V 3
G68V LPR 0:1
G68V LPR 4:1
G68V LPR 8:1
G68V LPR 16:1
G68V LPR 32:1
G68V LPR 60:1
G73A 1
G73A 2
G73A 3
G73A LPR 0:1
G73A LPR 4:1
G73A LPR 8:1
G73A LPR 16:1
G73A LPR 32:1
G73A LPR 60:1
G84A 1
G84A 2
G84A 3
G84A LPR 0:1
G84A LPR 4:1
G84A LPR 8:1
G84A LPR 16:1
G84A LPR 32:1
G84A LPR 60:1

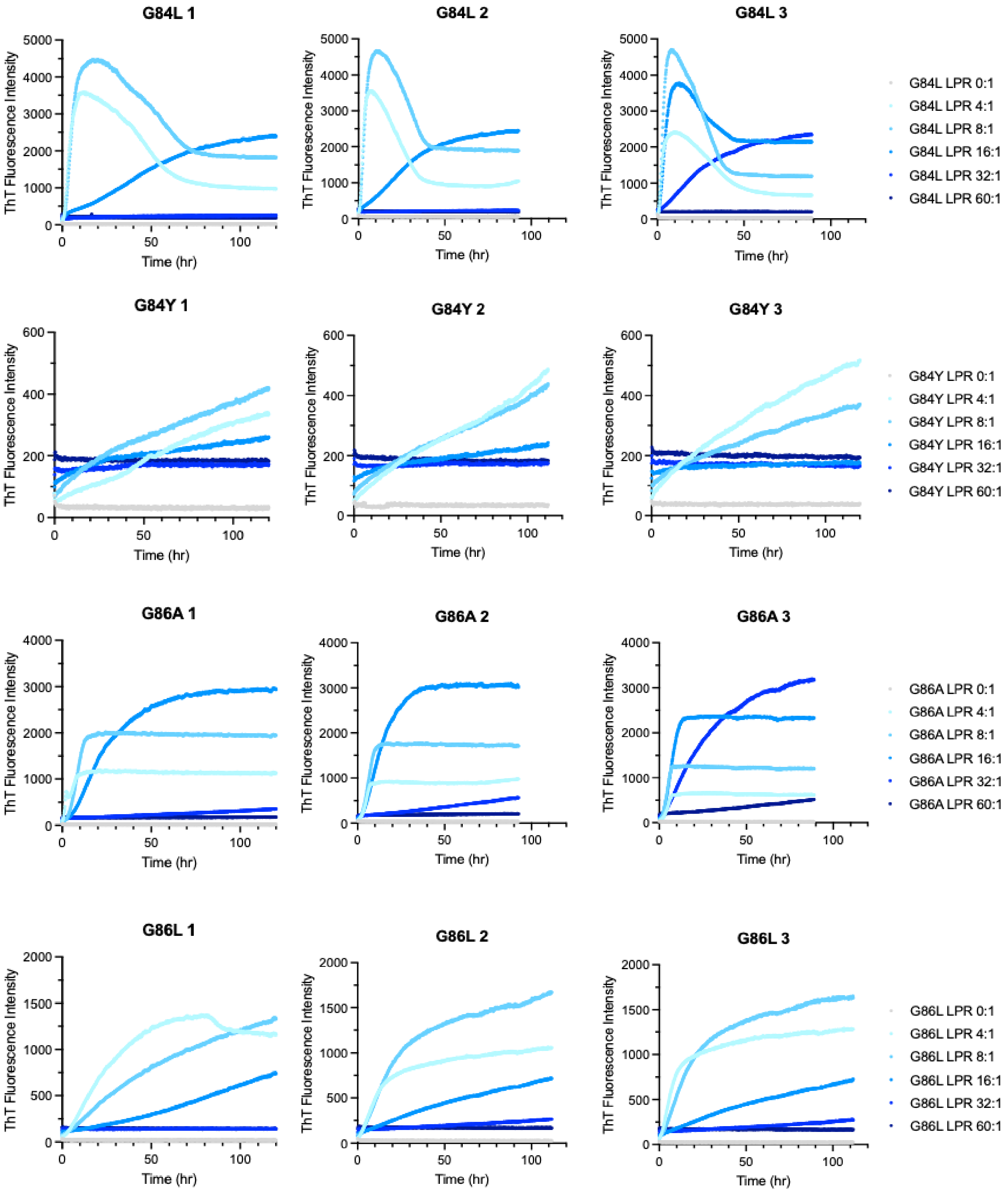


**Figure S5**. Thioflavin-T aggregation kinetics of α-synuclein variants at different ratios of DMPS lipid-to-protein ratios.

**(A)**

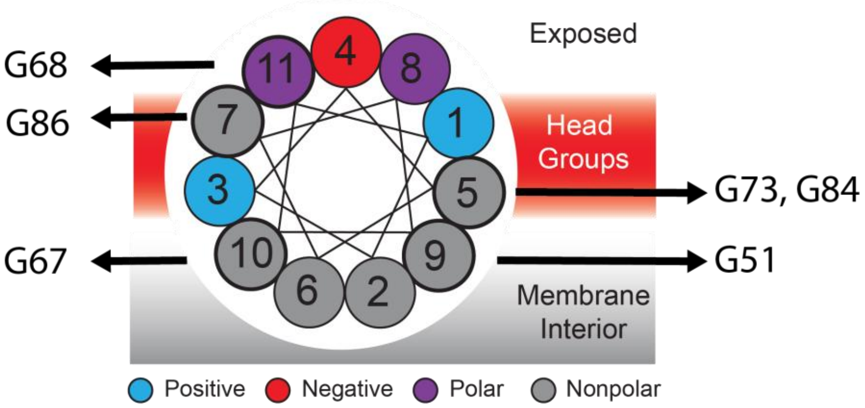


**(B)**

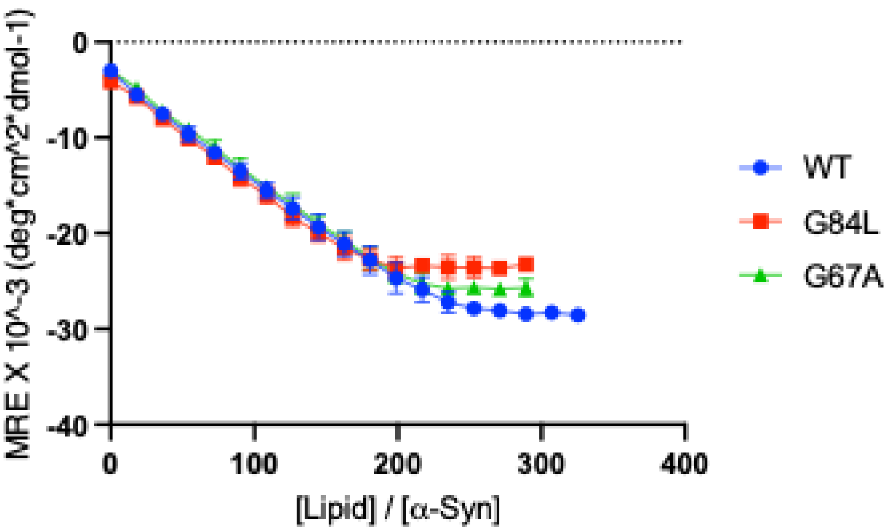


**Figure S6**. (A) Selected α-synuclein residues with the membrane surface. (B) Circular Dichroism Titration of purified α-synuclein WT and two non-polar variants at glycine positions with DMPS lipid vesicles ([α-synuclein] = 350 nM).

### Tables

**Table S1.** Aggregation scores and p-values of selected nonpolar substitutions at glycine residues in the NAC region.

| Variant | Aggregation score | p-value |
|---|---|---|
| **G51A** | 0.98401237 | 5.5974E-05 |
| **G67A** | 0.48413759 | 3.7492E-09 |
| **G67V** | 0.59996237 | 0.00035791 |
| **G68A** | 0.29872321 | 0.02801499 |
| **G68V** | 0.13038321 | 0.25929433 |
| **G73A** | 0.67796229 | 0.0671721 |
| **G84A** | 1.21724201 | 7.199E-05 |
| **G84L** | 1.19699804 | 1.2344E-26 |
| **G84Y** | 1.31220207 | 1.8727E-41 |
| **G86A** | 0.22840224 | 0.71799661 |
| **G86L** | -0.2133667 | 0.72365531 |

**Table S2.** List of DNA sequences.

| | |
|---|---|
| PCR 1 Forward Primer | ACACTCTTTCCCTACACGACGCTCTTCCGATCTACTCACTAT<br>AGGGAATATTAAGCTTGGTACCCTCGAG |
| PCR 1 Reverse Primer | GTGACTGGAGTTCAGACGTGTGCTCTTCCGATCTGGAAAG<br>CCAGTACCGATTTCTGCCAT |
| PCR 2 Forward Primer – UDI013 | AATGATACGGCGACCACCGAGATCTACACAAGGATGAACA<br>CTCTTTCCCTACACGACGC |
| PCR 2 Reverse Primer – UDI013 | GCACACGTCTGAACTCCAGTCACCCAAGTCTATCTCGTATG<br>CCGTCTTCTGCTTG |
| PCR 2 Forward Primer – UDI014 | AATGATACGGCGACCACCGAGATCTACACGGAAGCAGACA<br>CTCTTTCCCTACACGACGC |
| PCR 2 Reverse Primer – UDI014 | CAAGCAGAAGACGGCATACGAGATGAGTCCAAGTGACTGG<br>AGTTCAGACGTGTGC |
| Plasmid Sequence | CTGCATTAATGAATCGGCCAACGCGCGGGGAGAGGCGGTT<br>TGCGTATTGGGCGCTCTTCCGCTTCCTCGCTCACTGACTCG<br>CTGCGCTCGGTCGTTCGGCTGCGGCGAGCGGTATCAGCTC<br>ACTCAAAGGCGGTAATACGGTTATCCACAGAATCAGGGGAT<br>AACGCAGGAAAGAACATGTGAGCAAAAGGCCAGCAAAAG<br>CCCAGGAACCGTAAAAAGGCCGCGTTGCTGGCGTTTTTCC<br>ATAGGCTCCGCCCCCCTGACGAGCATCACAAAAATCGACG<br>CTCAAGTCAGAGGTGGCGAAACCCGACAGGACTATAAAGA<br>TACCAGGCGTTTCCCCCTGGAAGCTCCCTCGTGCGCTCTCC<br>TGTTCCGACCCTGCCGCTTACCGGATACCTGTCCGCCTTTC<br>TCCCTTCGGGAAGCGTGGCGCTTTCTCATAGCTCACGCTGT |

| | |
|---|---|
| | AGGTATCTCAGTTCGGTGTAGGTCGTTCGCTCCAAGCTGGG<br>CTGTGTGCACGAACCCCCCGTTCAGCCCGACCGCTGCGCC<br>TTATCCGGTAACTATCGTCTTGAGTCCAACCCGGTAAGACA<br>CGACTTATCGCCACTGGCAGCAGCCACTGGTAACAGGATTA<br>GCAGAGCGAGGTATGTAGGCGGTGCTACAGAGTTCTTGAA<br>GTGGTGGCCTAACTACGGCTACACTAGAAGGACAGTATTTG<br>GTATCTGCGCTCTGCTGAAGCCAGTTACCTTCGGAAAAAG<br>AGTTGGTAGCTCTTGATCCGGCAAACAAACCACCGCTGGT<br>AGCGGTGGTTTTTTTGTTTGCAAGCAGCAGATTACGCGCAG<br>AAAAAAAGGATCTCAAGAAGATCCTTTGATCTTTTCTACGG<br>GGTCTGACGCTCAGTGGAACGAAAACTCACGTTAAGGGAT<br>TTTGGTCATGAGATTATCAAAAAGGATCTTCACCTAGATCCT<br>TTTAAATTAAAAATGAAGTTTTAAATCAATCTAAAGTATATA<br>TGAGTAAACTTGGTCTGACAGTTACCAATGCTTAATCAGTG<br>AGGCACCTATCTCAGCGATCTGTCTATTTCGTTCATCCATAG<br>TTGCCTGACTCCCCGTCGTGTAGATAACTACGATACGGGAG<br>CGCTTACCATCTGGCCCCAGTGCTGCAATGATACCGCGAGA<br>CCCACGCTCACCGGCTCCAGATTTATCAGCAATAAACCAGC<br>CAGCCGGAAGGGCCGAGCGCAGAAGTGGTCCTGCAACTTT<br>ATCCGCCTCCATTCAGTCTATTAATTGTTGCCGGGAAGCTAG<br>AGTAAGTAGTTCGCCAGTTAATAGTTTGCGCAACGTTGTTG<br>GCATTGCTACAGGCATCGTGGTGTCACTCTCGTCGTTTGGT<br>ATGGCTTCATTCAGCTCCGGTTCCCAACGATCAAGGCGAGT<br>TACATGATCCCCCATGTTGTGCAAAAAAGCGGTTAGCTCCT<br>TCGGTCCTCCGATCGTTGTCAGAAGTAAGTTGGCCGCAGTG<br>TTATCACTCATGGTTATGGCAGCACTGCATAATTCTCTTACT<br>GTCATGCCATCCGTAAGATGCTTTTCTGTGACTGGTGAGTA<br>CTCAACCAAGTCATTCTGAGAATAGTGTATGCGGCGACCGA<br>GTTGCTCTTGCCCGGCGTCAATACGGGATAATAGTGTATCA<br>CATAGCAGAACTTTAAAAGTGCTCATCATTGGAAAACGTTC<br>TTCGGGGCGAAAACTCTCAAGGATCTTACCGCTGTTGAGAT<br>CCAGTTCGATGTAACCCACTCGTGCACCCAACTGATCTTCA<br>GCATCTTTTACTTTCACCAGCGTTTCTGGGTGAGCAAAAAC<br>AGGAAGGCAAAATGCCGCAAAAAAGGGAATAAGGGCGAC<br>ACGGAAATGTTGAATACTCATACTCTTCCTTTTTCAATGGGT<br>AATAACTGATATAATTAAATTGAAGCTCTAATTTGTGAGTTT<br>AGTATACATGCATTTACTTATAATACAGTTTTTTAGTTTTGCT<br>GGCCGCATCTTCTCAAATATGCTTCCCAGCCTGCTTTTCTGT<br>AACGTTCACCCTCTACCTTAGCATCCCTTCCCTTTGCAAATA<br>GTCCTCTTCCAACAATAATAATGTCAGATCCTGTAGAGACC<br>ACATCATCCACGGTTCTATACTGTTGACCCAATGCGTCTCCC<br>TTGTCATCTAAACCCACACCGGGTGTCATAATCAACCAATC<br>GTAACCTTCATCTCTTCCACCCATGTCTCTTTGAGCAATAAA<br>GCCGATAACAAAATCTTTGTCGCTCTTCGCAATGTCAACAG<br>TACCCTTAGTATATTCTCCAGTAGATAGGGAGCCCTTGCATG<br>ACAATTCTGCTAACATCAAAAGGCCTCTAGGTTCCTTTGTT<br>ACTTCTTCTGCCGCCTGCTTCAAACCGCTAACAATACCTGG<br>GCCCACCACACCGTGTGCATTCGTAATGTCTGCCCATTCTG<br>CTATTCTGTATACACCCGCAGAGTACTGCAATTTGACTGTAT<br>TACCAATGTCAGCAAATTTTCTGTCTTCGAAGAGTAAAAAA<br>TTGTACTTGGCGGATAATGCCTTTAGCGGCTTAACTGTGCC |

| | |
|---|---|
| | CTCCATGGAAAAATCAGTCAAGATATCCACATGTGTTTTTA<br>GTAAACAAATTTTGGGACCTAATGCTTCAACTAACTCCAGT<br>AATTCCTTGGTGGTACGAACATCCAATGAAGCACACAAGTT<br>TGTTTGCTTTTCGTGCATGATATTAAATAGCTTGGCAGCAAC<br>AGGACTAGGATGAGTAGCAGCACGTTCCTTATATGTAGCTT<br>TCGACATGATTTATCTTCGTTTCCTGCAGGTTTTTGTTCTGT<br>GCAGTTGGGTTAAGAATACTGGGCAATTTCATGTTTCTTCA<br>ACACTACATATGCGTATATATACCAATCTAAGTCTGTGCTCC<br>TTCCTTCGTTCTTCCTTCTGTTCGGAGATTACCGAATCAAAA<br>AAATTTCAAAGAAACCGAAATCAAAAAAAAGAATAAAAA<br>AAAAATGATGAATTGAATTGAAAAGCTAGCTTATCGATGAT<br>AAGCTGTCAAAGATGAGAATTAATTCCACGGACTATAGACT<br>ATACTAGATACTCCGTCTACTGTACGATACACTTCCGCTCAG<br>GTCCTTGTCCTTTAACGAGGCCTTACCACTCTTTTGTTACTC<br>TATTGATCCAGCTCAGCAAAGGCAGTGTGATCTAAGATTCT<br>ATCTTCGCGATGTAGTAAAACTAGCTAGACCGAGAAAGAG<br>ACTAGAAATGCAAAAGGCACTTCTACAATGGCTGCCATCAT<br>TATTATCCGATGTGACGCTGCAGCTTCTCAATGATATTCGAA<br>TACGCTTTGAGGAGATACAGCCTAATATCCGACAAACTGTT<br>TTACAGATTTACGATCGTACTTGTTACCCATCATTGAATTTT<br>GAACATCCGAACCTGGGAGTTTTCCCTGAAACAGATAGTAT<br>ATTTGAACCTGTATAATAATATATAGTCTAGCGCTTTACGGA<br>AGACAATGTATGTATTTCGGTTCCTGGAGAAACTATTGCATC<br>TATTGCATAGGTAATCTTGCACGTCGCATCCCCGGTTCATTT<br>TCTGCGTTTCCATCTTGCACTTCAATAGCATATCTTTGTTAA<br>CGAAGCATCTGTGCTTCATTTTGTAGAACAAAAATGCAACG<br>CGAGAGCGCTAATTTTTCAAACAAAGAATCTGAGCTGCATT<br>TTTACAGAACAGAAATGCAACGCGAAAGCGCTATTTTACCA<br>ACGAAGAATCTGTGCTTCATTTTTGTAAAACAAAAATGCAA<br>CGCGACGAGAGCGCTAATTTTTCAAACAAAGAATCTGAGC<br>TGCATTTTTACAGAACAGAAATGCAACGCGAGAGCGCTATT<br>TTACCAACAAAGAATCTATACTTCTTTTTTGTTCTACAAAAA<br>TGCATCCCGAGAGCGCTATTTTTCTAACAAAGCATCTTAGAT<br>TACTTTTTTTCTCCTTTGTGCGCTCTATAATGCAGTCTCTTG<br>ATAACTTTTTGCACTGTAGGTCCGTTAAGGTTAGAAGAAGG<br>CTACTTTGGTGTCTATTTTCTCTTCCATAAAAAAAGCCTGAC<br>TCCACTTCCCGCGTTTACTGATTACTAGCGAAGCTGCGGGT<br>GCATTTTTTCAAGATAAAGGCATCCCCGATTATATTCTATAC<br>CGATGTGGATTGCGCATACTTTGTGAACAGAAAGTGATAGC<br>GTTGATGATTCTTCATTGGTCAGAAAATTATGAACGGTTTCT<br>TCTATTTTGTCTCTATATACTACGTATAGGAAATGTTTACATT<br>TTCGTATTGTTTTCGATTCACTCTATGAATAGTTCTTACTACA<br>ATTTTTTTGTCTAAAGAGTAATACTAGAGATAAACATAAAA<br>AATGTAGAGGTCGAGTTTAGATGCAAGTTCAAGGAGCGAA<br>AGGTGGATGGGTAGGTTATATAGGGATATAGCACAGAGATA<br>TATAGCAAAGAGATACTTTTGAGCAATGTTTGTGGAAGCGG<br>TATTCGCAATGGGAAGCTCCACCCCGGTTGATAATCAGAAA<br>AGCCCCAAAAACAGGAAGATTGTATAAGCAAATATTTAAAT<br>TGTAAACGTTAATATTTTGTTAAAATTCGCGTTAAATTTTTG<br>TTAAATCAGCTCATTTTTTAACGAATAGCCCGAAATCGGCA<br>AAATCCCTTATAAATCAAAAGAATAGACCGAGATAGGGTTG |

| | |
|---|---|
| | AGTGTTGTTCCAGTTTCCAACAAGAGTCCACTATTAAAGAA<br>CGTGGACTCCAACGTCAAAGGGCGAAAAAGGGTCTATCAG<br>GGCGATGGCCCACTACGTGAACCATCACCCTAATCAAGTTT<br>TTTGGGGTCGAGGTGCCGTAAAGCAGTAAATCGGAAGGGT<br>AAACGGATGCCCCCATTTAGAGCTTGACGGGGAAAGCCGG<br>CGAACGTGGCGAGAAAGGAAGGGAAGAAAGCGAAAGGA<br>GCGGGGGCTAGGGCGGTGGGAAGTGTAGGGGTCACGCTG<br>GGCGTAACCACCACACCCGCCGCGCTTAATGGGGCGCTAC<br>AGGGCGCGTGGGGATGATCCACTAGTACGGATTAGAAGCC<br>GCCGAGCGGGTGACAGCCCTCCGAAGGAAGACTCTCCTCC<br>GTGCGTCCTCGTCCTCACCGGTCGCGTTCCTGAAACGCAG<br>ATGTGCCTCGCGCCGCACTGCTCCGAACAATAAAGATTCTA<br>CAATACTAGCTTTTATGGTTATGAAGAGGAAAAATTGGCAG<br>TAACCTGGCCCCACAAACCTTCAAATGAACGAATCAAATTA<br>ACAACCATAGGATGATAATGCGATTAGTTTTTTAGCCTTATT<br>TCTGGGGTAATTAATCAGCGAAGCGATGATTTTTGATCTATT<br>AACAGATATATAAATGCAAAAACTGCATAACCACTTTAACT<br>AATACTTTCAACATTTTCGGTTTGTATTACTTCTTATTCAAAT<br>GTAATAAAAGTATCAACAAAAAATTGTTAATATACCTCTATA<br>CTTTAACGTCAAGGAGAAAAAACCCCGGATCGGACTACTA<br>GCAGCTGTAATACGACTCACTATAGGGAATATTAAGCTTGG<br>TACCCTCGAGatggatgtattcatgaaaggactttcaaaggccaaggagggagttgtgg<br>ctgctgctgagaaaaccaaacagggtgtggcagaagcagcaggaaagacaaaagagggtgtt<br>ctctatgtaggctccaaaaccaaggagggagtggtgcatggtgtggcaacagtggctgagaag<br>accaaagagcaagtgacaaatgttggaggagcagtggtgacgggtgtgacagcagtagccca<br>gaagacagtggagggagcagggagcattgcagcagccactggctttgtcaaaaaggaccagtt<br>gggcaagaatgaagaaggagccccacaggaaggaattctggaagatatgcctgtggatcctga<br>caatgaggcttatgaaatgccttctgaggaagggtatcaagactacgaacctgaagccATGG<br>CAGAAATCGGTACTGGCTTTCCATTCGACCCCCATTATGTG<br>GAAGTCCTGGGCGAGCGCATGCACTACGTCGATGTTGGTC<br>CGCGCGATGGCACCCCTGTGCTGTTCCTGCACGGTAACCCG<br>ACCTCCTCCTACGTGTGGCGCAACATCATCCCGCATGTTGC<br>ACCGACCCATCGCTGCATTGCTCCAGACCTGATCGGTATGG<br>GCAAATCCGACAAACCAGACCTGGGTTATTTCTTCGACGAC<br>CACGTCCGCTTCATGGATGCCTTCATCGAAGCCCTGGGTCT<br>GGAAGAGGTCGTCCTGGTCATTCACGACTGGGGCTCCGCT<br>CTGGGTTTCCACTGGGCCAAGCGCAATCCAGAGCGCGTCA<br>AAGGTATTGCATTTATGGAGTTCATCCGCCCTATCCCGACCT<br>GGGACGAATGGCCAGAATTTGCCCGCGAGACCTTCCAGGC<br>CTTCCGCACCACCGACGTCGGCCGCAAGCTGATCATCGATC<br>AGAACGTTTTTATCGAGGGTACGCTGCCGATGGGTGTCGTC<br>CGCCCGCTGACTGAAGTCGAGATGGACCATTACCGCGAGC<br>CGTTCCTGAATCCTGTTGACCGCGAGCCACTGTGGCGCTTC<br>CCAAACGAGCTGCCAATCGCCGGTGAGCCAGCGAACATCG<br>TCGCGCTGGTCGAAGAATACATGGACTGGCTGCACCAGTC<br>CCCTGTCCCGAAGCTGCTGTTCTGGGGCACCCCAGGCGTT<br>CTGATCCCACCGGCCGAAGCCGCTCGCCTGGCCAAAAGCC<br>TGCCTAACTGCAAGGCTGTGGACATCGGCCCGGGTCTGAA<br>TCTGCTGCAAGAAGACAACCCGGACCTGATCGGCAGCGAG<br>ATCGCGCGCTGGCTGTCGACGCTCGAGATTTCCGGCTAANN<br>NNNNNNNNNNNNNNNATGGTCCTGCTGGAGTTCGTGACCGC |

| | |
|---|---|
| | CGCCGGGATCACTCTCGGCATGGACGAGCTGTACAAGTAA<br>GATCCTCGAGCATGCATCTAGAGGGCCGCATCATGTAATTA<br>GTTATGTCACGCTTACATTCACGCCCTCCCCCCACATCCGCT<br>CTAACCGAAAAGGAAGGAGTTAGACAACCTGAAGTCTAGG<br>TCCCTATTTATTTTTTTATAGTTATGTTAGTATTAAGAACGTT<br>ATTTATATTTCAAATTTTTCTTTTTTTTTCTGTACAGACGCGTG<br>TACGCATGTAACATTATACTGAAAACCTTGCTTGAGAAGGT<br>TTTGGGACGCTCGAAGGCTTTAATTTGCGGCC |
